# Top-down prefrontal connectivity accompanies perceptual breakthrough after interocular suppression

**DOI:** 10.64898/2026.11.12.765589

**Authors:** Kavindu H. Bandara, Elise G. Rowe, Olivia Carter, Marta I. Garrido

**Affiliations:** Melbourne School of Psychological Sciences, The University of Melbourne, Parkville, Victoria, Australia; Graeme Clark Institute for Biomedical Engineering, The University of Melbourne, Australia

## Abstract

The role of the prefrontal cortex (PFC) in the emergence of visual stimuli into conscious awareness remains contested. Additionally, whether top-down prefrontal connectivity is shaped by prior expectations during this process is not well understood. To address these questions, we applied dynamic causal modelling (DCM) to an EEG dataset during breaking continuous flash suppression (bCFS), which offers tight control over when a stimulus breaks into conscious perception. Our network comprised bilateral PFC, primary visual cortex, fusiform gyrus and motor cortex. Because perceptual breakthrough precedes behavioural report, we used a temporally resolved classifier to empirically define each participant’s moment of perceptual breakthrough (*M* = 1250ms, *SD* = 100ms). DCMs were then inverted over this time window up until each individual’s breakthrough point. Parametric empirical Bayes revealed that top-down connectivity from the PFC to all regions was necessary to explain the observed neural dynamics (posterior probability [pp] > 0.99), though evidence that this top-down influence was modulated by expectation was inconclusive (pp = 0.74). Together, these findings suggest that top-down PFC connectivity accompanies perceptual breakthrough in bCFS, but it remains unclear whether this is modulated by expectation.

## Introduction

The precise role of the prefrontal cortex (PFC) during the emergence of a conscious percept remains an open question, with direct implications for its broader role in consciousness. At the core of this debate are the neural mechanisms governing how the brain resolves sensory ambiguity during interocular competition. One view proposes that this competition is actively resolved in frontal areas (Brascamp et al., 2018; Kapoor et al., 2022; Logothetis et al., 1996; Lumer et al., 1998; Panagiotaropoulos et al., 2012; Weilnhammer et al., 2013), while the other argues that this resolution occurs more locally within visual areas (Blake, 1989; Haynes et al., 2005; Hesse & Tsao, 2020; Polonsky et al., 2000; Sheinberg & Logothetis, 1997; Tong et al., 1998; Tong & Engel, 2001). One potential avenue to make progress in this literature would be to use paradigms where a stimulus gradually enters into awareness.

Broadly, interocular suppression paradigms that evoke perceptual transitions of stimuli into and out of awareness have been particularly valuable to understand the neural processes governing how a stimulus overcomes suppression and gains conscious access. In this phenomenon, a stimulus presented to one eye is rendered temporarily invisible by competing input to the other eye, allowing researchers to track the conditions under which suppressed information gains entry to awareness. The most widely studied example is binocular rivalry, where distinct images are presented dichoptically and the resulting percept oscillates stochastically between the two monocular inputs as they vie for dominance (Blake & Logothetis, 2002). However, during binocular rivalry, perceptual transitions occur spontaneously and unpredictably, and its phenomenology can be quite complex; sometimes involving gradual, piecemeal transitions rather than discrete switches between stable percepts (Yilmaz et al., 2025). This can make isolating the neural dynamics during the moment of perceptual entry rather ambiguous.

Instead, paradigms such as breaking continuous flash suppression (bCFS; Jiang et al., 2007; Stein et al., 2011; Tsuchiya & Koch, 2005) provides a key methodological advantage for assessing the influence of frontal regions as a stimulus enters awareness. bCFS uses a high-contrast dynamic mask to initially suppress a target stimulus until it overcomes interocular suppression and eventually *breaks through* into conscious awareness. This design reliably evokes awareness of the suppressed stimulus on every trial, making it well-suited for investigating whether top-down frontal connectivity accompanies the emergence of a conscious percept.

As the speed of breakthrough in bCFS can be used to infer the prioritisation of certain types of stimuli to enter awareness, the current study additionally interrogated how frontal contributions might be modulated by the manipulation of expectation. It has been shown that prior expectations modulate the threshold and speed at which stimuli enter into conscious perception (Meijs et al., 2018; Melloni et al., 2011; Pinto et al., 2015); though the direction of the effect may depend on contextual factors (Press et al., 2020). Frontal effects are of particular interest here because the influence of expectation necessarily requires that high-level semantic information reaches higher-order cortical regions. As such, top-down connectivity may suppress early sensory regions for expected stimuli and releases these regions from suppression when expectations are violated; consistent with a predictive processing account of perceptual inference (Friston, 2005; Rao & Ballard, 1999; Summerfield & de Lange, 2014).

Although bCFS offers methodological advantages for studying neural dynamics as suppressed stimuli gain conscious access, neuroimaging and electrophysiological evidence specific to this paradigm remains sparse. To our knowledge, only a handful of studies have examined the neural correlates of breakthrough during bCFS (McFadyen et al., 2022; del Río et al., 2018; Travis, 2020; Kalhan et al., 2022). Of these studies, only Kalhan et al. (2022) directly examined the role of frontal areas during bCFS, demonstrating frontoparietal and occipital activation correlated with faster evidence accumulation during bCFS. However, as fMRI has limited temporal resolution, this study offered limited insight into the directed connectivity arising from frontal networks as stimuli break through into awareness. Here, we leverage bCFS and temporally resolved EEG data to investigate the directed influence of prefrontal regions during perceptual breakthrough.

Recent work from our lab employed a bCFS paradigm to examine how prior expectations and affective valence influence the speed at which suppressed stimuli breakthrough into awareness (McFadyen et al., 2022). Crucially, however, this previous study did not examine connectivity between higher-order and sensory regions during these breakthrough events, leaving open the question of whether there is a directed influence of frontal areas to lower regions during the emergence of visual stimuli into awareness. To investigate this, we use a technique known as dynamic causal modelling (DCM) for evoked responses. DCM estimates directed effective connectivity between neural regions by inverting a generative model of the observed scalp data (David et al., 2006). We hypothesised that if frontal regions are involved in conscious emergence during bCFS, then top-down connectivity from the PFC will be necessary to explain the neural dynamics during the breakthrough window. If these predictions are confirmed by the data, we further hypothesise that prior expectations will modulate this top-down connectivity from the PFC during the breakthrough window, with stronger top-down connectivity predicted for expected compared to unexpected stimuli.

## Methods

### Participants

The current study is a secondary analysis of an EEG dataset previously acquired by our lab (McFadyen et al., 2022). The original sample consisted of 33 healthy adults. For the present analysis, three participants were excluded: one for having an excessive non-response rate (41.4% of all trials), and a further two due to technical errors with their EEG triggers. Thus, the final sample taken to the analysis stage consisted of 30 participants (14 males, 16 females, *M* = 22 years old, SD = 2.1 years).

### Experimental task

Whilst comprehensive experimental details are available in the original publication (McFadyen et al., 2022), a summary of the task is provided here for context (see Figure 1). This experiment incorporated a bCFS paradigm to investigate how prior expectations and affective stimuli were prioritised during visual processing. The frequency of fearful versus neutral faces was manipulated across counterbalanced blocks to create facial emotions that were either expected or unexpected. Within a given block, the frequent facial emotion (the expected stimulus) appeared on 83% of trials, while the infrequent emotion (the unexpected stimulus) appeared on 17% of trials. In each trial, a flashing Mondrian mask (10Hz) was presented to one monocular channel, whilst a face stimulus was presented to the other. The contrast of the face stimulus increased linearly in contrast over a 3-second window, peaking at a titrated, participant-specific maximum to ensure conscious breakthrough thresholds were roughly equal across the cohort. Participants were instructed to indicate the orientation of the face (tilted 5° clockwise or counter-clockwise) via a button press the moment the face broke through into awareness and were instructed to prioritise both speed and accuracy.

**Figure 1.**
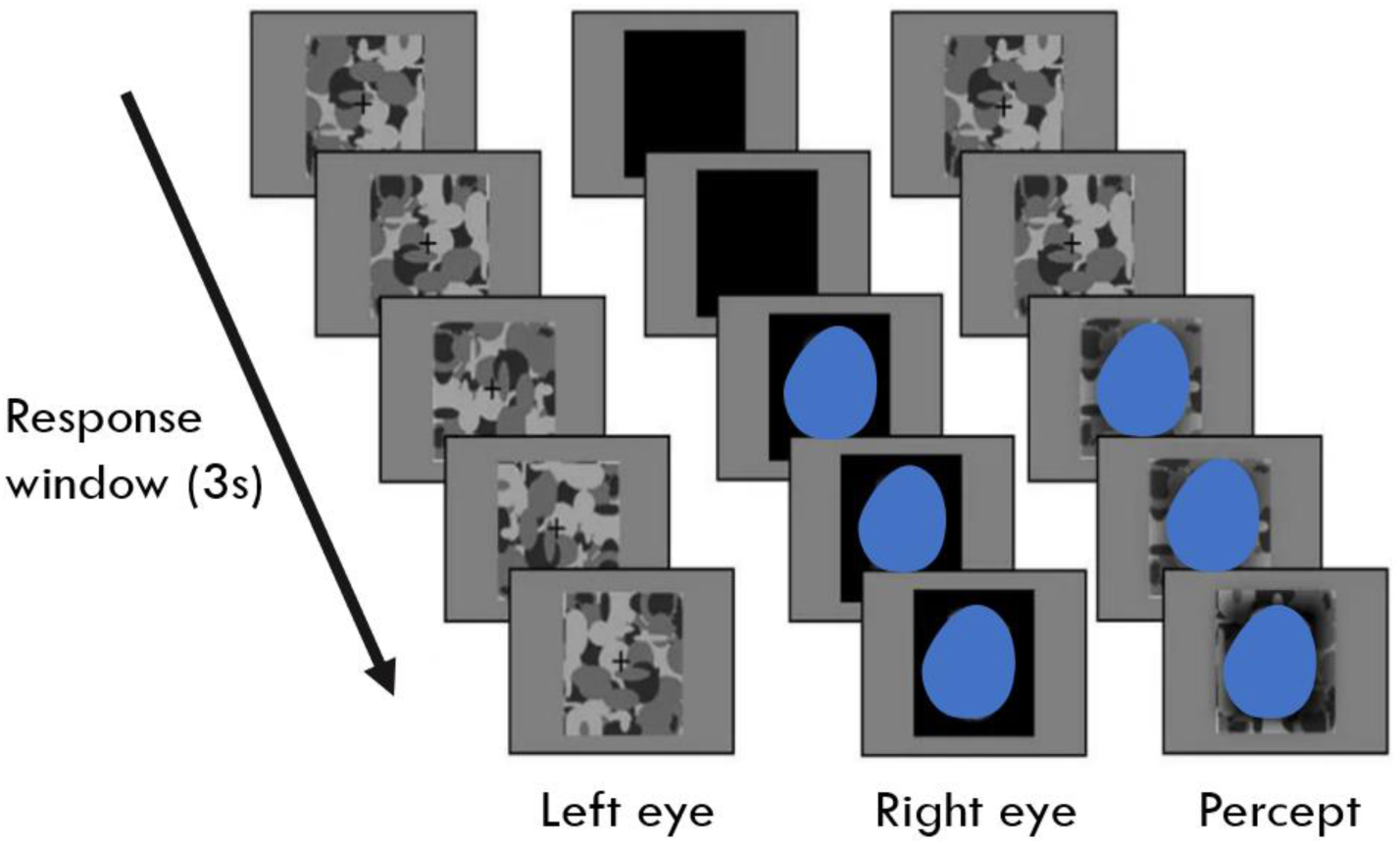
Schematic of the breaking Continuous Flash Suppression (bCFS) paradigm adapted from McFadyen et al. (2022). A standard trial unfolded over a 3-second response window. Distinct visual streams were presented dichoptically (that is, a different image was shown to each eye). A high-contrast, 10Hz flickering mask (left column) was presented to one eye, initially suppressing awareness of the target stimulus, a face (middle column), presented to the contralateral eye. As the trial progressed, the contrast of the target face gradually increases. The resulting perceptual experience (approximated in the right column) demonstrates the target eventually overcoming interocular suppression and breaking through into conscious awareness. Note, the real face images used in this study were replaced by blue ovals to comply with bioRxiv’s policy to avoid identifying information of people. See McFadyen et al. (2022; Figure 1B) for the original illustration.

### EEG Preprocessing

Continuous 64-channel EEG data were preprocessed using a combination of MNE-Python (v1.8.0; Gramfort et al., 2014) and SPM12 (https://www.fil.ion.ucl.ac.uk/spm/) in MATLAB 2024b (The MathWorks Inc., Natick, MA). This pipeline combined MNE-Python for automated independent components analysis (ICA) artifact rejection with SPM12 for robust averaging, source reconstruction, and DCM. All data was pre-processed and analysed on the high-performance computing platform, Spartan, at the University of Melbourne.

Prior to ICA, the EEG data was downsampled to 200Hz and high-pass filtered at 1Hz. ICA is typically done on a downsampled and filtered version of the data to ensure ICA stability and data quality (Winkler et al., 2015), and then applied to the original data. Independent components were decomposed using the infomax algorithm (Lee et al., 1999), retaining sufficient components to explain 99% of the signal variance per participant. The ICLabel classifier (implemented in MNE-python; Li et al., 2022; Pion-Tonachini et al., 2019) was used to automatically detect artefactual components at a confidence threshold of 95%. An average of 4.55 components (SD = 3.34) were identified and subtracted from the continuous EEG data.

The cleaned data was then converted into an SPM readable format and the remainder of preprocessing was conducted in SPM. To attenuate line noise and high-frequency interference, a band-pass filter (0.1-40 Hz) and a notch filter (49-51 Hz) were applied. To remove mask related flickering at 10Hz (and its harmonics), a 200ms moving average filter was also applied. Artifact rejection was performed using a thresholding approach, where channel amplitude exceeding 100 µV were marked as bad, and segments containing more than 20% bad channels were subsequently rejected. Epochs were defined from −100ms to 3000ms relative to stimulus onset and each epoch was labelled according to the four trial types – either expected fearful, unexpected fearful, expected neutral, or unexpected neutral. Trials were excluded if a participant did not respond, responded incorrectly, or if the response was faster than 500ms. These latter trials were interpreted as indicating an accidental button press, as the face contrast at this time was very low and it was unlikely participants were aware of the face before this timepoint. Per participant, a mean of 69.5 (5.52%) trials were marked for having an incorrect or no response, and 8.2 trials were marked for premature responses (0.65%). Accounting for overlap between these criteria, a total of 73.2 (5.82%) unique trials were excluded per participant, such that on average a total of 1150 trials (*SD* = 81 trials) were brought to the analysis stage.

The preprocessing pipeline diverged at this point for data that underwent multivariate decoding, and data that was source reconstructed and underwent DCM analysis. This was because decoding requires single-trial data whereas the DCM data handles averaged evoked response data well. For multivariate decoding analysis, the artifact-corrected data were baseline-corrected (again from −100ms to 0ms) without robust averaging. To reduce motor contamination during the training window, trials were further excluded if a response occurred after 2700ms only for the decoding dataset. For source reconstruction and DCM analysis, this data was subjected to robust averaging. A secondary low-pass filter at 40 Hz was applied to remove high-frequency noise introduced by the averaging process (Litvak et al., 2011). Trials were then baseline-corrected from −100ms to 0ms.

### Identifying individual temporal windows of interest

Given the face stimulus in the bCFS paradigm gradually ramped in contrast, typical visual evoked potentials were not observed in this dataset (i.e, no distinct N170 component is observable; McFadyen et al., 2022). Consequently, we adopted a data-driven multivariate approach to empirically define the temporal windows of interest for the subsequent DCM.

We employed a support vector machine (SVM) algorithm with a radial basis function (RBF) kernel to characterise the neural transition of the face as it breaks into awareness. The classifier was trained to distinguish between two unambiguous perceptual states. The early time window (which we will henceforth refer to as the *mask window*) spanned from 200-500ms. This window captured visual processing dominated by the flashing mask. We omitted the first 200ms during training to avoid the high-amplitude, evoked response at the very start of the trial induced by the flickering mask. Up until 500ms, the face was at an extremely low contrast and it was reasoned that participants’ experience would consist of only the mask and not of the face. This was well before the mean response time at 1810ms. The late time window (which we will refer to hereafter as the *face window*) spanned from 2700ms to 3000ms (the end of the trial). This window corresponded to the period of maximal face contrast, post-response, when the participant had explicitly indicated that they were fully aware of the stimulus. Here, the aim was to identify the 50% point between the mask and face, such that this served as each participant’s individual breakthrough point.

As shown in figure 2A, a 7-fold cross-validation scheme was implemented across the trial dimension. In each fold, the classifier was trained on the defined mask and face windows from six blocks and tested on the remaining block. To prevent data leakage, features (i.e. channel voltages) were normalised (z-scored) based on the training set statistics, and then these parameters subsequently applied to the test set (Lemm et al., 2011; Varoquaux et al., 2017).

**Figure 2.**
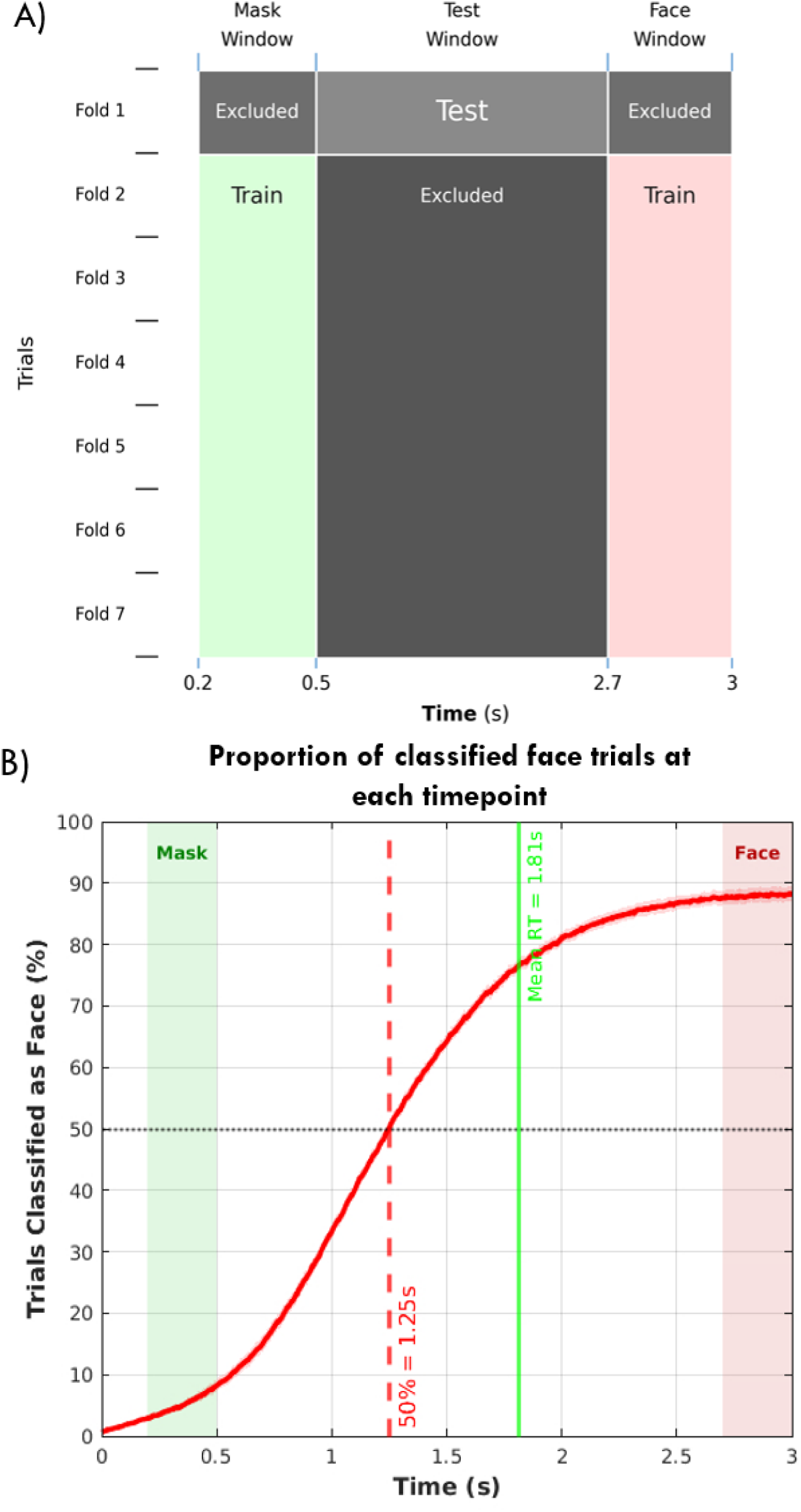
Data-driven identification of DCM analysis time window using a multivariate pattern classifier. **A)** A schematic of the 7-fold cross-validation scheme applied across trials. A support vector machine (SVM) classifier was trained to distinguish between two unambiguous perceptual states: an early mask-dominated window (0.2-0.5s) and a late face-dominated window (2.7-3.0 s). For each fold, the classifier was trained on six blocks and tested on the remaining block. To generate a time-resolved prediction of the neural state, the trained classifier was applied to the ambiguous transition window (0.5–2.7 s). **B)** The proportion of trials classified as face-like at each timepoint across the epoch. The green and red shaded regions represent the mask and face training windows, respectively. The solid red curve illustrates the time-resolved prediction of the neural state, which was calculated as the proportion of trials belonging to one of the two classes predicted by the SVM at each timestep. The vertical dashed red line marks the 50% crossover threshold (mean = 1.25 s). This threshold serves as an empirical marker indicating when the scalp topography reliably departed from the mask window, and it was used to define the end of the time window for the subsequent DCM analysis on an individual basis. The solid green line indicates the mean behavioural response time at 1.81s.

The trained classifier was then applied to the ambiguous transition window (500-2700ms) to generate a time-resolved prediction of the neural state; that is, whether the scalp topography at each timepoint more closely resembled the face window or the mask window. Raw SVM scores (distance from the decision boundary) were binarized at each time point (corresponding to 5ms per timestep as the EEG data was downsampled to 200Hz), providing a discrete prediction of whether the SVM classified each time point as mask-like or face-like. We then computed the proportion of trials which were identified as either mask-like or face-like across the epoch.

To determine the precise window for the DCM analysis, we sought to identify a time window of when the perceptual breakthrough likely occurred on an individual basis. We evaluated this multivariate evidence alongside previous univariate EEG and behavioural analyses of this dataset; specifically, change point analyses (which identify statistical shifts in single-channel EEG activity) and drift diffusion modelling of behavioural reaction times previously conducted by McFadyen et al. (2022) to estimate the onset of evidence accumulation during decision-making processes). In that paper, a late change point in the EEG data (occurring at 1137ms) was found to correlate strongly with sensory evidence accumulation (drift rate), while the period thereafter correlated with post-decision processing parameters. However, this correlational analysis was restricted to a subsample of participants (20 out of the 31), whereas the present multivariate procedure was applied to the full analytic sample, potentially lending to a more statistically stable estimate. Additionally, because multivariate decoding evaluates the distributed spatial pattern across all electrodes simultaneously, it is inherently more sensitive to widespread neural changes rather than single-channel amplitude shifts (Hebart & Baker, 2018), affording greater sensitivity compared to the previous change-point procedure.

We identified the point at which the decoder transitioned from more mask-like to face-like (or the 50% threshold) and used this time point as the end of the DCM time window on an individual basis (see Figure 2B). It is important to clarify how the 50% crossover is interpreted here. The crossover point is not interpreted as the onset of perceptual breakthrough nor does it *necessarily* index the end of evidence accumulation. Rather, it reflects the timepoint at which the majority of trials have transitioned to a neural response more similar to the face window than the mask window. It is therefore best understood as an empirical, data-driven marker of when the scalp topography has reliably departed from the mask window. Accordingly, the mean 50% threshold (occurring at 1250ms) fell within a broadly comparable range to the independent estimate of the end of sensory evidence accumulation reported by McFadyen et al. (2022) as 1137ms on average across participant, albeit using a different method on a partially overlapping subsample. This 113ms difference may be non-trivial, given the rapidity of visual processing and because the two approaches are methodologically quite distinct (we discuss this in more detail in the Discussion). Nevertheless, the similarity suggests the estimate is of the right order of magnitude. On this basis, we defined the window of interest from 500ms until the 50% threshold; an interval assumed to span the perceptual breakthrough process, and importantly, ended well before the mean response time (1810ms), thereby minimising motor execution confounds. The time period post 500ms was chosen because the contrast of the face prior to 500ms was presumed to be too low to be visible to participants. Hence starting the modelling window from 500ms aims to capture dynamics during the time window in which the target face emerges into awareness.

### Identifying the nodes for DCM

To define the nodes for the DCM analysis, we adopted a hybrid approach of using a data-driven approach that was restricted over certain regions based on our research question of interest. Specifically, the EEG data were source reconstructed and anatomical masks were then applied to extract region-specific peak coordinates. We performed a group-level inversion across all 30 participants, a procedure that constrains the source solution to be consistent across subjects, thereby improving the accuracy of source estimates (Litvak & Friston, 2008).

A canonical cortical mesh was used to define the source space, with coregistration performed using standard fiducials from the SPM anatomical template. The forward model was computed using an EEG boundary element model. Source inversion was achieved using the multiple sparse priors algorithm with a greedy search scheme (Friston et al., 2008). The inversion was constrained to a time window of 500 to 1250ms relative to stimulus onset, and spanned the full frequency range (0–256 Hz). The resulting source images were smoothed to 12mm^3^.

Regions of Interest (ROIs) were defined using anatomical masks generated from the JuBrain Anatomy Toolbox (Eickhoff et al., 2005). Masks were created for the bilateral primary visual cortex (V1; Amunts et al., 2000), fusiform gyrus (FG; Caspers et al., 2013; Lorenz et al., 2017), motor cortex (M1; Geyer et al., 1996), and PFC. For the PFC mask, we included all frontal areas (Amunts et al., 1999; Amunts et al., 2004; Bludau et al., 2014; Henssen et al., 2016; Geyer, 2012; Geyer et al., 1996) but explicitly excluded the supplementary motor area, to avoid conflating frontal contributions with premotor activity.

To identify the dipole locations for the DCM, a 2 (emotion) × 2 (expectation) full-factorial general linear model (GLM) was applied to the source images. To capture the perceptual switch common to all trial types, we computed the mean effect across all four conditions (i.e. a contrast vector of: “[1 1 1 1]”). The peak MNI coordinates for this contrast were extracted within each anatomical mask and subsequently used as the prior source locations for the DCM analysis.

### DCM Model Specification

Effective connectivity was estimated using DCM for evoked responses (David et al., 2006). We employed an equivalent current dipole spatial model with an ERP neuronal model (Jansen & Rit, 1995). Given that the experiment involved constantly flickering stimuli alongside a face which ramped up in contrast, we shifted the onset of the modelling time window to start at 500ms (instead of 0 ms) until the MVPA derived 50% crossover threshold (defined per individual; noting the mean time was 1250ms across the sample). This window aimed to capture the dynamics associated with breakthrough of the face. To determine the most appropriate model, we inverted four different DCMs per participant and chose the best model via subsequent Bayesian model selection. Here, we sought to determine what architecture (either including or not including a motor node) and input function (either a discrete, transient input or continuous, gradual input) characterised the observed data the best (as described in more detail below).

First, we tested two distinct network architectures, either a 6-node network comprising the bilateral V1, FG, and PFC nodes or an 8-node network with additional bilateral M1 nodes (see Figure 3A). While the motor cortex was not a region of theoretical interest here (our hypotheses concern top-down connectivity from the PFC to lower areas), preparatory motor activity could plausibly occur within the analysis window and influence the estimation of PFC connectivity. Although the mean end point of the analysis window (1250ms) terminated well before the mean response time (1810ms; thereby minimising confounds arising from motor *execution*) its timing does not preclude the presence of motor *preparation* within the window. There is substantial evidence that preparatory motor signals emerge considerably in advance of overt responding: the readiness potential arises approximately 1.5 s prior to movement onset (Shibasaki & Hallett, 2006) and, critically, activity in motor cortex builds up gradually during evidence accumulation during perceptual decision-making several seconds before the manual report in some paradigms (Donner et al., 2009; Lange et al., 2013). It is therefore plausible that such preparatory activity may fall within the window analysed here. Were this activity present but unmodelled, the remaining sources could have absorbed its contribution to the scalp signal, confounding the connectivity estimates of interest. We therefore modelled bilateral M1 as a source, so that motor-related variance would be attributed to a dedicated motor node and top-down frontal connectivity could be left to reflect putative prefrontal, rather than motor-related, contributions. We therefore did not assume the presence or absence of a motor contribution, but specified it as a candidate node and tested its necessity empirically.

**Figure 3.**
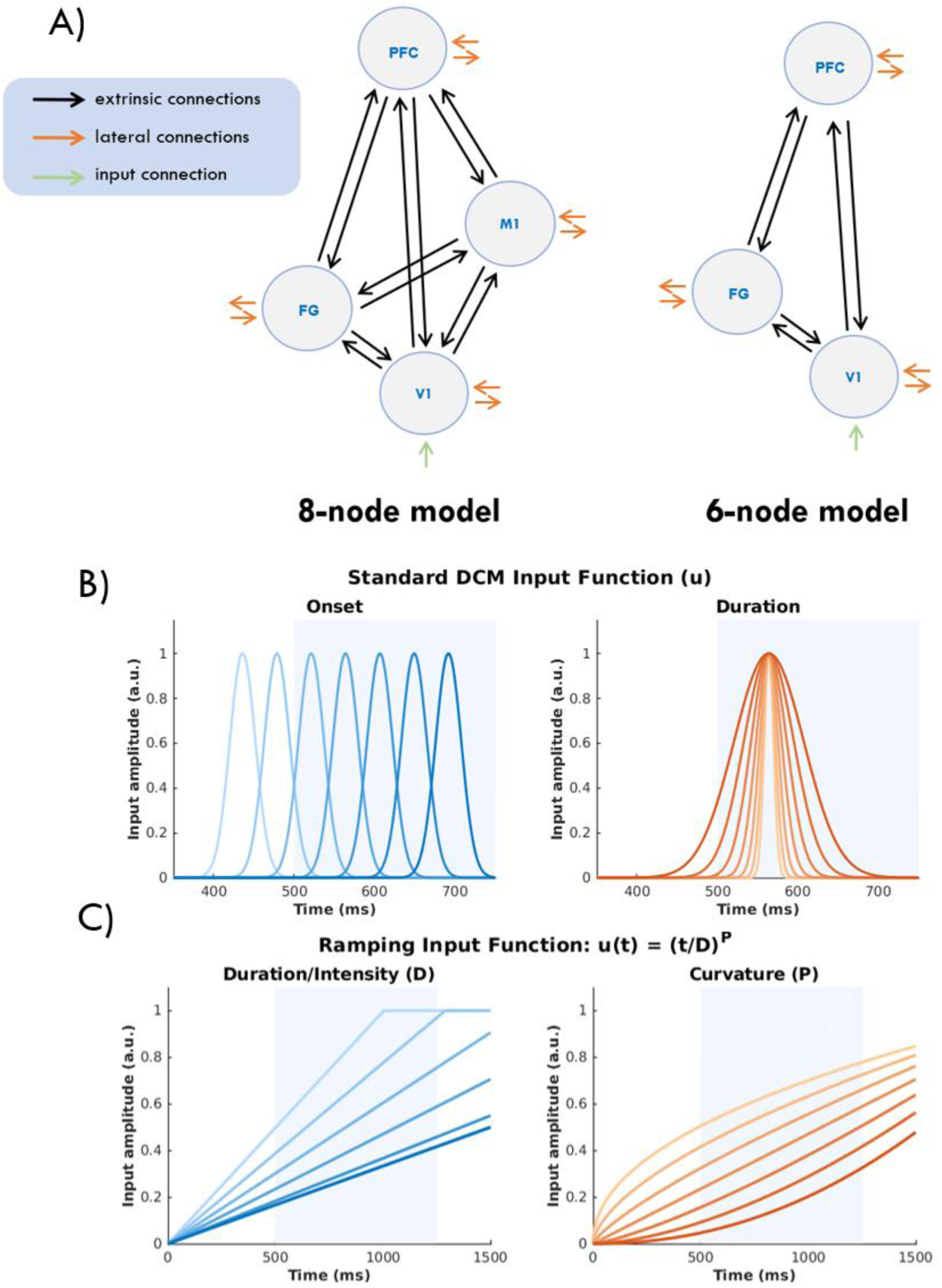
The four different DCM model types. This shows both the model architectures and the two input functions. **(A)** Network architectures for the 8-node (left) and 6-node (right) models. Nodes represent the bilateral primary visual cortex (V1), fusiform gyrus (FG), prefrontal cortex (PFC), and motor cortex (M1; present in the 8-node model only). This diagram only shows one hemisphere, but both hemispheres were modelled. Green arrows show the sensory input, black arrows indicate extrinsic (forward and backward) connections, while orange arrows represent lateral connections between hemispheres. The panels illustrate the range of input profiles available to the DCM inversion scheme. **(B)** One of the two input functions tested. To illustrate how these free parameters change the resulting input function, we show how each free parameter might vary when the other free parameter is fixed at a certain value. In practice, however, both parameters are jointly optimised for each participant when the model is inverted. For the standard input function, this is a conventional discrete (Gaussian) input typically employed in DCM for evoked responses. The curves represent the different values of stimulus onset (when width is fixed at its prior value of 16ms) and temporal dispersion (when onset is fixed at its prior value of 64ms post-window onset). In all plots, the shaded blue region denotes the mean time window selected used in the DCM analysis (500-1250ms). **(C)** The second input function tested, the ramping input function. This was defined as 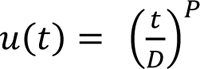 and this function models the gradual accumulation of sensory information. The left panel demonstrates variations in intensity/slope (parameter *D*) under a linear assumption (*P* = 1). The right panel depicts variations in curvature (parameter *P*) when the duration parameter, D, is fixed at its prior value, 2125ms, which is the midpoint between the mean of the MVPA-defined crossover point across participants (1250ms) and the duration of the stimulus (3000ms).

The candidate network architectures were fully connected at each level and visual input was specified at the bilateral V1 nodes (the C-matrix). We allowed all intrinsic and extrinsic connections (forward, backward, and lateral) to be modulated by the experimental conditions of interest (the B-matrix). Here, we modelled the modulatory effect of only expectation (using a contrast vector of [−1 1 −1 1]) because the main effect of emotion and their interaction were not of theoretical interest.

We tested two different input functions for our DCMs that modelled afferent thalamic input, *u(t)*, as either a discrete input event or a gradual increase. These models were then formally compared using Bayesian model selection (BMS). This approach was used as it may not have been biologically accurate to use the standard DCM input for this type of paradigm because the stimuli was not a brief transient presentation and instead ramped up over time. Thus, we sought to establish whether matching the input to the visual properties of the paradigm more closely might fit the observed data better.

The first input function was the standard DCM input function for evoked responses. Formally, this input is defined as a Gamma density function; however, under standard experimental conditions, this is approximated as a Gaussian (David et al., 2006). This function represents a discrete, time-locked afferent volley (originating from the thalamus), where the shape reflects the temporal dispersion of presynaptic action potentials due to axonal conduction delays and synaptic integration times (David et al., 2006). In the standard DCM implementation, the onset time is a free parameter with a prior (or default value) of 64ms post-stimulus, corresponding to the conduction delay from the retina to the early visual cortex via the thalamic input. A second free parameter controls the width (dispersion) of this volley (see Figure 3B). However, as we are modelling the sensory input of a stimulus (the face) that gradually increases in contrast, it was hypothesised that this experimental data may not be best captured by a discrete, phasic onset of a standard evoked response. Additionally, because the dataset did not exhibit a typical N170 component, this suggested that the underlying neural driver of the facial input might not be best characterised by a standard discrete input. Within the DCM framework, model inference is always relative to the model space provided. Thus, there is a risk that the *true* generative process lies outside the tested hypotheses. Therefore, to explicitly test whether the sensory drive during bCFS is better characterised as a continuous accumulation rather than a discrete event, we specified a second input model and formally compared this input against the standard input function by comparing their model evidence (free energy) via BMS. This allowed us to determine whether the data were better explained by a transient afferent volley or a gradual accumulation.

To explicitly test whether the input to V1 might be gradual, the alternative input model was based on Stevens’ power law (Stevens, 1957). We selected this power law function because it captured the gradual ramping relationship between physical stimulus magnitude, *I*, and perceived intensity, ψ, governed by the equation, ψ ∝ *I^k^*. Additionally, Stevens’ law is a relatively simple, psychophysical law that has been found to characterise perception across a wide array of sensory modalities, demonstrating that subjective intensity consistently scales as a power function of physical magnitude; with the precise power depending on the exact stimulus (Stevens, 1957; 1961). Treating perceived intensity as the relevant input to early visual cortex, this framework naturally motivates a power-law input function for the present paradigm, where the physical contrast ramp is expected to drive a non-linearly scaled neural response.

Crucially, while the ramping of the stimulus in our design was linear, we allowed a non-linear component in the function to approximate the transduction properties of the early visual cortex (Albrecht & Hamilton, 1982). That is, neural firing rates in V1 do not map linearly to input contrast; but rather follow a (sigmoidal) contrast response function. Although a full sigmoidal input would be more biologically precise, this would require *two* free parameters (slope and half-maximal contrast) to estimate the curvature alone (before adding in a third parameter for the duration). To match the number of free parameters across both input functions (so both contain two) and thus ensure that any differences between the input functions are attributable to differences in the temporal profile of the inputs (rather than model complexity), we opted for a single parameter to approximate the curvature component. This allows the inversion to empirically estimate whether the breakthrough into awareness is driven by a non-linear accumulation of sensory evidence, while avoiding over-parameterising the model.

We further modified the canonical power law equation, ψ ∝ *I^k^*, because the neuronal state equations in DCM require inputs to be bounded within a physiologically plausible dynamic range (typically normalized to unity). An unbounded power function with an arbitrary scaling coefficient could grow indefinitely with time, destabilising the ordinary differential equations governing the neural mass model. Hence, the scaling coefficient was re-parameterized by introducing a duration parameter, *D*, to express the input as normalised over elapsed time, bounding the function such that *u*(*t*) = 1 when t = D (that is, it saturates at maximal values of D). A prior of 2125ms was chosen for this duration parameter as it was the midpoint between the mean crossover point (at 1250ms, representing the average end point of the analysed DCM window, and hence the point expected to be the earliest time the duration parameter reaches its maximum contrast) and the end of the trial (representing the last timepoint the duration reaches its maximum contrast). This only serves as a prior value, and is a free parameter which is optimised during model fitting. Since the stimulus contrast in the present design increases linearly with time, *t* was substituted for physical intensity, *I*, to map the temporal contrast increase to the neural input function. This yields:

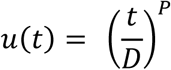

where *P* governs the curvature of the function and *D* sets the timescale over which the input reaches its maximum. The onset was fixed at 0ms to coincide with the physical start of the trial, such that all variance in the input profile is explained by the curvature (*P*) and duration (*D*) parameters alone (see Figure 3C).

For completeness, the standard Gaussian input (approximating a Gamma density), representing a discrete, transient event is:

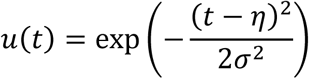

where *η* (onset latency) and *σ* (width) are free parameters controlling the timing and dispersion of the afferent thalamic volley.

Finally, as a quality check, we assessed model performance by computing the mean variance explained across all DCMs in all four types of models tested. This was calculated 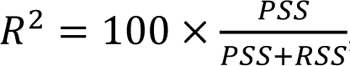, where, the predicted sum of squares (PSS) represents the signal energy captured by the model (*PSS = ∑_i_∑_j_H^2^_ij_*) and the residual sum of squares (RSS) represents the unexplained variance (*RSS = ∑_i_∑_j_R^2^_ij_*), summed over time bins, *i*, and channels, *j*.

### Identifying the network and input function

To determine the optimal network architecture and input function, we inverted the four types of DCMs per participant and formally compared these using random-effects (RFX) BMS in a 2 (network architecture: 8-node vs 6-node DCM) x 2 (input type: standard vs ramping) factorial model space (Stephan et al., 2009). Inference was performed at the family level (Penny et al., 2010), pooling evidence for all models sharing a specific feature (e.g., the ramping input family versus the Gaussian input family).

To quantify the confidence in our model selection, we report the protected exceedance probability (PEP; Rigoux et al., 2014). Unlike standard exceedance probability, which describes the probability that a model is more likely than any other, PEP additionally corrects for the null hypothesis that observed differences in model evidence are due to chance rather than reflecting a true effect. A PEP > 0.95 indicates strong evidence that the winning family is the most prevalent model in the population, above and beyond chance.

### Parametric empirical Bayes

Following BMS, we used parametric empirical Bayes (PEB) to estimate group-level connectivity parameters for the winning model architecture. PEB implements a hierarchical Bayesian framework in which random effects on connection strengths are modelled explicitly, effectively treating individual subject parameters as samples from a group-level distribution (Friston et al., 2016). Within this framework, we tested two specific hypotheses regarding the role of the prefrontal cortex (PFC) using Bayesian model reduction (BMR) and Bayesian model comparison (BMC) as follows.

First, we tested whether top-down frontal connectivity from the PFC was necessary to explain the average observed neural dynamics across all conditions (i.e. the A-matrix). We compared the evidence of the full model (containing all specified connections) against a reduced model where bilateral PFC backward connections in the A-matrix were removed by fixing their prior mean to zero. As A-matrix parameters represent the mean effective connectivity across the experimental session, this test determined whether a baseline top-down signal is required to generate the data within the specified time window of interest.

Second, we tested whether the main effect of expectation modulated this top-down influence. Here, we modelled expectation as a modulatory input on the B-matrix. We compared the full model (where expectation was free to modulate all connections) against a reduced model where the modulatory parameter for PFC backward connections to all areas were removed.

For both tests, the posterior probability (pp) of the model was calculated from the difference in log-model evidence (approximated by the change in free energy, Δ*F*) between the full and reduced models using 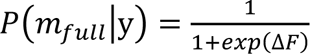. Here, a pp > 0.75 constitutes positive evidence and a pp > 0.95 constitutes strong evidence for the necessity of the connection or modulation (Kass & Raftery, 1995).

## Results

BMS analysis of different DCM architectures revealed that a network containing motor nodes were necessary to explain the data, and in addition, the winning model also favoured a gradual ramping input over the standard discrete volley used in DCM. Consistent with our primary hypothesis, DCM revealed that top-down connectivity from the PFC was necessary to explain the neural data during the breakthrough window. However, contrary to our secondary predictions, this top-down connectivity was not modulated by prior expectations. We detail the specific results below.

### Nodes identified for DCM

The group-level source inversion yielded a mean explained variance of 89.51% (SD = 11.81%), indicating that the source model captured the observed scalp topography with a high degree of fidelity. The peak MNI coordinates (shown in Figure 4) extracted from the GLM analysis of the mean effect of all conditions within each anatomical mask were as follows: left V1 [−2, −92, −4], right V1 [16, −100, −4], left fusiform gyrus [−42, −44, −20], right fusiform gyrus [40, −46, −22], left M1 [−6, −20, 62], right M1 [12, −24, 70], left PFC [−6, 62, 16], and right PFC [6, 62, 12]. The resulting locations served as prior dipole positions for all subsequent DCM analyses.

**Figure 4.**
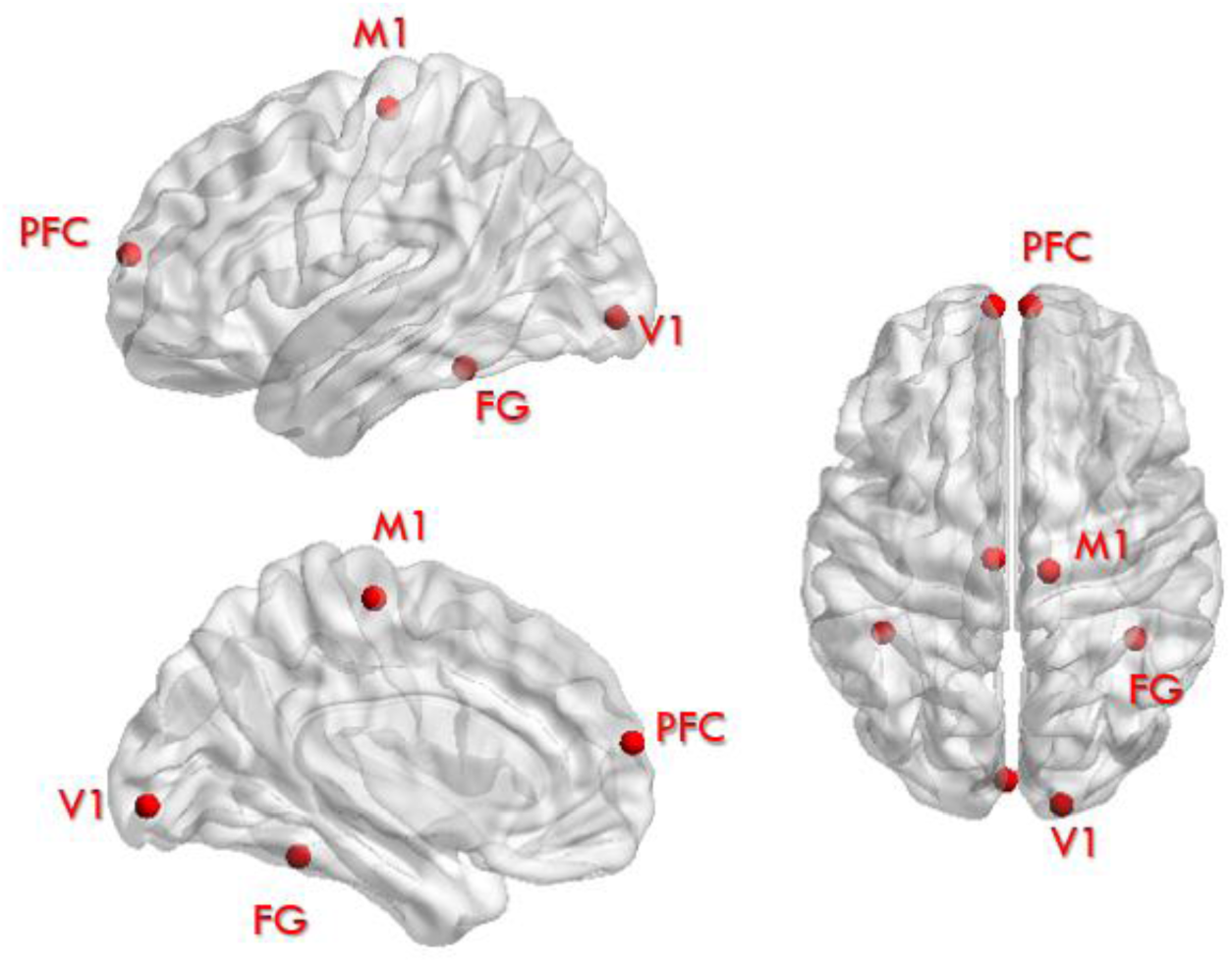
Source priors for dynamic causal modelling. Left lateral, left medial, and superior views of the group-level peak coordinates from the mean effect of all conditions from the GLM analysis, superimposed on a transparent standard MNI template brain. Red markers indicate the dipole locations used for the DCM. PFC: prefrontal cortex, M1: motor area, FG: fusiform gyrus, V1: primary visual cortex.

#### Network architecture and input function comparison

To adjudicate between the competing hypotheses regarding input dynamics and network architecture, we employed RFX BMS. We first performed family-level inference in a 2 (input: standard vs ramping) x 2 (6-node vs 8-node) factorial design (with and without M1). The fitted input functions for each model are shown in Figure 5A.

**Figure 5.**
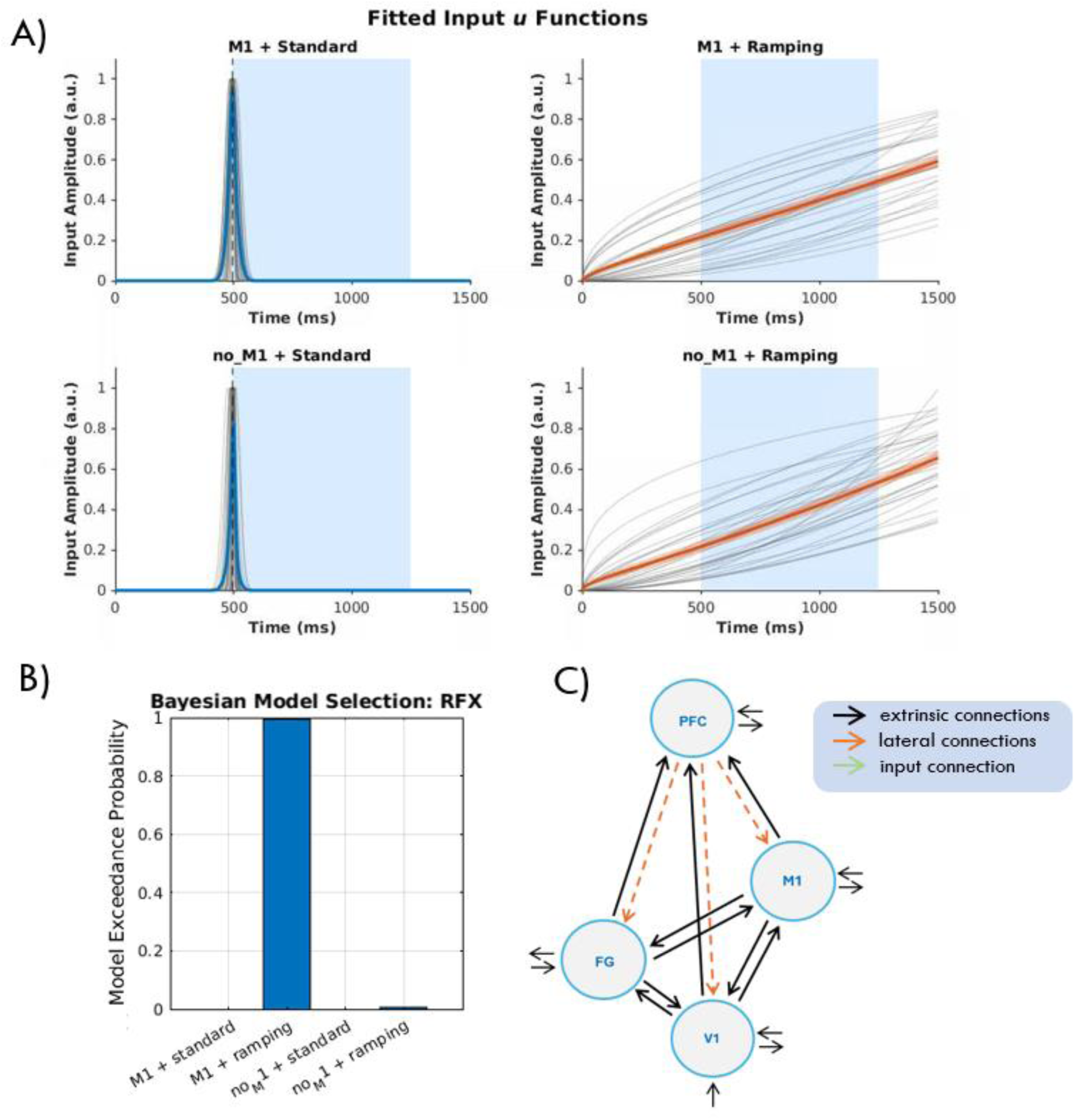
Fitted input functions and results from Bayesian model selection for each of the four DCM architectures tested and network tested via PEB. **A)** The left side shows the standard (Gaussian) input functions, shown for the M1 (top) and No-M1 (bottom) architectures. The fitted Gaussians are centred around the onset of the analysis window at 500ms (left) and the ramping (power law) input functions for the corresponding architectures (right). Individual participant functions (grey) reveal variability in the fitted curvature parameter *P*, with some participants exhibiting convex (accelerating) and others concave (decelerating) accumulation profiles. In all panels, the group mean is shown in blue (standard) or orange (ramping), with the shaded band denoting ±1 standard error of the mean, and grey lines indicating fitted functions at the participant-level. The blue shaded region denotes the mean DCM analysis window (500–1250ms). **B)** Results of the random-effects (RFX) Bayesian Model Selection (BMS). This shows the model-level BMS across all four individual models. **C)** Architecture of the reduced parametric empirical Bayes (PEB) model used in the A- and B-matrix tests. Nodes represent the primary visual cortex (V1), fusiform gyrus (FG), motor cortex (M1), and prefrontal cortex (PFC). Solid black arrows denote forward and lateral connections allowed to vary, whilst dashed orange arrows denote the PFC connections that were fixed to a value of zero (switched off).

As a quality check to ensure that the specified DCMs tested provided a reasonable representation of the observed signals, we computed the accuracy of the model fit as the variance explained by the model. The model inversion yielded a satisfactory goodness-of-fit, with a mean variance explained of 95.24% (SD = 3.57%) averaged across participants for the winning model. All model variance explained is shown in Table 1. This confirms that the DCM, especially the novel ramping input model, successfully captured the spatiotemporal dynamics of the scalp topography. Note that this only serves as a check to ensure the models were capturing variance in the data (especially the ramping models, as this was a novel approach here), and should not be used for comparative purposes given the difference in the number of parameters between the models. Hence, we use BMS for formal comparison of the different model types (see below).

**Table 1.** Mean variance explained and standard deviation for each of the four DCM architectures tested, computed as the proportion of scalp signal energy captured by the model averaged across all participants. All four models achieved satisfactory goodness-of-fit. *denotes the winning model as identified by Bayesian model selection.

| **Model** | **Mean Variance Explained** | **SD** |
|---|---|---|
| M1 + Ramping* | 95.24% | 3.57% |
| M1 + Standard | 92.08% | 5.60% |
| No M1 + Ramping | 93.80% | 5.75% |
| No M1 + Standard | 92.78% | 4.74% |

The first comparison assessed the validity of the input function. The family of models which used the power law approximation input demonstrated decisive evidence over the standard Gaussian input family (protected exceedance probability [PEP] > 0.999). This result empirically validates our modelling of the “breakthrough” of the face as a cumulative, non-linear process rather than a transient, stimulus-locked event. The second comparison examined the necessity of the motor nodes in the network architecture. The family of models including motor nodes (M1 models) outperformed the models without M1 (PEP > 0.999). This indicates that activity within the motor system is essential for explaining the variance in the EEG data during the perceptual transition. Finally, comparison of individual models identified the M1 with the bespoke ramping input architecture as the best model, with a PEP of 0.974.

### Testing the necessity of top-down frontal-posterior connectivity

Having established the winning model architecture, PEB was employed to test our two pre-specified hypotheses regarding firstly its necessity during perceptual breakthrough (see Figure 5C), and secondly its modulation by prior expectation. Comparison of the full model against a reduced model in which bilateral PFC backward connections were removed yielded a pp > 0.99, showing very strong evidence that top-down connectivity from the PFC is necessary during the breakthrough window.

The second analysis tested whether this top-down influence was dynamically modulated by prior expectation (via the B-matrix). Here, we observed ambiguous evidence that the modulatory effect of expectation on PFC backward connectivity was required to explain the data (pp = 0.74 in favour of a reduced model without B-matrix modulations). That is, whilst top-down frontal-posterior connectivity is robustly present in the mean connectivity across conditions (the A-matrix), we did not observe strong evidence suggesting that this connectivity is modulated according to whether the incoming stimulus was expected or unexpected.

## Discussion

Here, we investigated the role of the PFC during the emergence of a face into conscious awareness during bCFS. Using a MVPA decoding algorithm, we first identified a time window during which the EEG data switched from a mask-like to a face-like neural state on an individual basis. We then characterised the effective connectivity from prefrontal regions as suppressed stimuli emerged into awareness during this putative breakthrough window for each participant using DCM. In addition, we developed a novel ramping input function and demonstrated that it better explained the data compared to the standard discrete input function traditionally used in DCM for evoked responses. We found that top-down connectivity from the PFC to sensorimotor regions is necessary during perceptual breakthrough. However, we did not observe modulation in top-down connectivity from the PFC as a function of stimulus expectation over this time window. This was despite behavioural data demonstrating an effect of expectation on response times (McFadyen et al., 2022). Taken together, these findings align with the view that top-down frontal-posterior connectivity is important as stimuli reaches conscious during interocular suppression.

### Top-down PFC connectivity accompanies perceptual breakthrough

Here, we demonstrate that top-down frontal-posterior connectivity supports perceptual breakthrough during bCFS. This finding is in alignment with evidence implicating frontoparietal areas in resolving perceptual rivalry (Brascamp et al., 2018; Lumer et al., 1998; Lumer & Rees, 1999; Qian et al., 2026; Weilnhammer et al., 2021; Weilnhammer et al., 2013). Although neuroimaging studies during bCFS are relatively sparse compared to other forms of perceptual rivalry, the few studies that exist suggest a role of frontoparietal areas in bCFS, in addition to visual regions (del Río et al., 2018; Kalhan et al., 2022); though importantly, these studies did not examine connectivity. Addressing this gap, we demonstrate that top-down frontal connectivity accompanies the transition into awareness during bCFS within our defined breakthrough time window.

An important caveat to our findings is that whilst we observed top-down frontal connectivity during perceptual breakthrough, we cannot uniquely associate this PFC connectivity to a perceptual process. This limitation primarily relates to the lack of a baseline condition in this study (i.e. the stimulus goes from unseen to seen in all conditions), such that the underlying neural activity may likely be a mixture of perceptual, decision-making, motor-preparation, and other post-perceptual processes that occur in the vicinity of consciousness breakthrough. To more conclusively establish whether this top-down frontal connectivity is perceptual in nature, future work could aim to contrast these connectivity dynamics against a physically identical baseline condition under which the stimulus stays unseen. This could be achieved, for instance, by titrating the target contrast to increase and plateau at a participant’s individual threshold (rather than at full contrast) such that the stimuli remain physically matched across breakthrough and suppressed trials. Additionally, adapting the paradigm to remove the requirement for immediate report (i.e., a no-report paradigm) would further minimise the influence of post-perceptual confounds (we discuss this in more detail below).

An alternative interpretation of the current findings is that frontoparietal areas may not necessarily drive the emergence of a percept but rather select between competing representations that have already been resolved at lower levels of the visual hierarchy. Accordingly, we cannot rule out that early visual competition co-occurred in our data. Recent evidence from Qian et al. (2025) demonstrates that higher regions actively resolve perceptual ambiguity, even whilst early visual areas exhibit local interocular competition. Using ultra-high-field 7T fMRI, that study showed that although initial sensory conflict manifests locally within the superficial layers of the primary visual cortex, the resolution of this competition was heavily modulated by top-down connectivity from parietal regions (i.e. the intraparietal sulcus; IPS). Specifically, they demonstrated that switch-related activity in the IPS preceded activity in early visual areas and subsequent DCM analysis indicated an increase in top-down connectivity emanating from the IPS during rivalry. However, one alternative explanation for these findings is that competition may indeed still be resolved in other visual areas, such that the IPS they observed merely provides attentional boosts to expedite the transition; akin to demonstrations that top-down contextual cues accelerate local sensory processing (Bar, 2004). By extension, it is plausible that the top-down frontal connectivity observed in the current study may likewise reflect a selective or modulatory contribution which may expedite or bias a transition whose underlying competition is resolved within visual areas; rather than driving perceptual breakthrough per se. (Filimon et al., 2013)

Since participants were actively reporting the moment of breakthrough on every trial, we cannot rule out that top-down frontal connectivity observed here reflects, at least in part, the report-related and post-perceptual demands these studies associate with frontal engagement, rather than perceptual breakthrough per se. Indeed, some of the most pressing evidence against the frontal theories comes from experiments where reporting confounds are more strictly controlled. When perceptual transitions are rendered invisible or unreportable, transition-related responses in the frontoparietal network are largely eliminated (Frässle et al., 2014; Brascamp et al., 2015; Zou et al., 2016). Because interocular competition relies on endogenous changes in conscious perception, standard versions of these tasks frequently require observers to continuously report the nature of their percept. This reliance on active reporting in classic rivalry paradigms conflates the neural correlates of conscious perception with post-perceptual processes required for report (Aru et al., 2012). When addressing this confound, experiments which delay the time of report from the experience or use other objective^1^ metrics to infer awareness (termed no-report tasks; Tsuchiya et al., 2015) have challenged the view that frontal involvement is strictly perceptual and may instead reflect reporting requirements (although see Bandara et al., 2025; Hatamimajoumerd et al., 2022; Kapoor et al., 2022). For example, Frässle et al. (2014) used objective physiological measures (namely, the optokinetic nystagmus and pupil size) to infer and continuously track perceptual experience during report and no-report conditions. They demonstrated that when observers passively experienced rivalry, differential neural activity in frontal areas was absent, whereas activation in occipital and parietal regions persisted supporting the view that frontal areas are associated with post-perceptual demands rather than driving the rivalry itself. It remains an open question whether enhanced frontoparietal activity drives perceptual transitions, or whether it reflects downstream consequences of decision-making, task-monitoring, and other post-perceptual processes (Aru et al., 2012). While the present study elicited report during each trial, the development of a no-report variant of the bCFS task represents an important direction for future work. This would allow a direct test of whether top-down frontal connectivity observed here is genuinely perceptual or, contingent on the demands of report.

### A ramping input function better characterises gradual sensory stimulation

The present BMS findings suggest that sensory input should be parameterised differently in paradigms in which stimulation accumulates gradually rather than arriving as a rapid discrete event. In this regard, our ramping input function represents a novel methodological contribution with broader applicability. Whilst standard DCM for evoked responses uses a transient Gaussian function, appropriate for paradigms employing abrupt stimulus onsets, we demonstrate that this form of input is suboptimal for bCFS. Within the model space tested here, the ramping input function had the strongest evidence, indicating that the neural dynamics during bCFS are better characterised as continuous accumulation than as a transient, stimulus-locked event. Although paradigms lacking a discrete onset are sometimes analysed as experimental effects using DCM for the complex cross-spectra (Friston et al., 2012), that formulation of DCM is designed for steady-state data and rests on stationarity assumptions; namely, that the underlying neural dynamics remain statistically stable over the analysed time window. However, this assumption is often violated in task-evoked paradigms, such as bCFS, where the input to the network is non-stationary. More broadly, the same approach used here may be worth considering in other paradigms with gradual onset stimulation. A few examples might be in some sustained attention tasks (Fortenbaugh et al., 2017); naturalistic stimuli (Huk et al., 2018; Sonkusare et al., 2019); contrast-change evidence accumulation tasks (O’Connell et al., 2012; Steinemann et al., 2018); gradual onset continuous performance tasks (Esterman et al., 2013; Kucyi et al., 2020); and, visual (Franconeri & Simons, 2003) and auditory (Benghanem et al., 2024; Neuhoff, 1998) looming stimuli. Alternative input functions in such paradigms that more faithfully capture the temporal structure of the sensory stimulus may lead to more robust models

Additional limitations concerns the empirical definition of our analysis window. We utilised MVPA to identify the temporal onset of the perceptual transition; however, this involved several analytical choices that, while principled, may not have been purely determined by the data. Foremost among these is the use of a 50% crossover as the endpoint of each participant’s window. While this represents a natural midpoint and proxy for a temporal marker of perceptual breakthrough, a stricter or more lenient threshold, or different boundaries for the mask and face training windows may have shifted the window in time.. Though the similarity between our data-driven approach to define the breakthrough window (mean at 1250ms) and the previously reported estimate of the end of sensory evidence accumulation (at 1137ms; McFadyen et al., 2022) provides some reassurance, the temporal window identified by these two methods differed by approximately 113ms. This may be considered a non-trivial difference given the rapidity of visual perception. Additionally, these two estimates are derived from methodologically distinct approaches; targeting related, but not identical, constructs. Though their proximity is encouraging, it is possible that alternative windowing choices might yield different connectivity estimates and inferences.

A further complication arises from the nature of how MVPA itself works. Notably, the late training window (2.7-3s) was necessarily post-response, as this was the period in which face contrast was maximal and participants had confirmed awareness of the face. Accordingly, the classifier could have learned to discriminate a pre-perceptual state from a post-decisional one, meaning the crossover reflects a transition between these two states in addition to perceptual emergence of the face. A decoder will necessarily exploit *any* widespread co-occurring signals, including motor preparation and post-decisional processes. Insofar as motor preparatory and post-decisional signals generate distributed EEG patterns during the transition window, they may contribute to the classifier’s discrimination, and the direction of any resulting bias on the final estimate is difficult to determine a priori. We adopted this window on the assumption that perceptual breakthrough and the termination of evidence accumulation occur closely in time, an assumption supported by the convergence of our crossover with the independent estimate of the end of evidence accumulation reported by McFadyen et al. (2022). We therefore interpret the crossover as marking the point at which the scalp topography has reliably departed from the mask-dominated state, around which perceptual breakthrough is likely to occur. We nonetheless acknowledge that we cannot explicitly rule out some contribution of post-decisional processes within the analysed time window.

## Conclusions

In summary, this study provides evidence for top-down frontal connectivity to sensorimotor regions during perceptual breakthrough during bCFS. We also provide a methodological advance to modelling data from paradigms where the stimulus has a gradual-onset and which benefit from input functions that more faithfully capture the temporal structure of the stimulus. While we show evidence that frontal influences are needed here, future work should aim to develop no-report variants of this task to further characterise the specific contribution of the PFC to the emergence of conscious visual perception, independently of motor responses.

## Code availability

https://github.com/kbandara/bcfs

## Acknowledgements

This research was supported by The University of Melbourne’s Research Computing Services and the Petascale Campus Initiative. This research was supported by the Commonwealth through an Australian Government Research Training Program Scholarship (DOI: https://doi.org/10.82133/C42F-K220) and the Melbourne School of Psychological Sciences PhD Studentship.

## Footnotes

1 Objective in this sense refers to the fact that the measure used to infer conscious perception does not rely on introspective reports of one’s experience.

